# HOPE-SIM, a cryo-structured illumination fluorescence microscopy system for accurately targeted cryo-electron tomography

**DOI:** 10.1101/2022.09.16.508243

**Authors:** Shuoguo Li, Xing Jia, Tongxin Niu, Xiaoyun Zhang, Chen Qi, Wei Xu, Hongyu Deng, Fei Sun, Gang Ji

## Abstract

Cryo-focused ion beam (cryo-FIB) milling technology has been developed for the fabrication of cryo-lamella of frozen native specimens for study by *in situ* cryo-electron tomography (cryo-ET). However, the precision of the target of interest is still one of the major bottlenecks limiting application. Here, we developed a new cryo-correlative light and electron microscopy (cryo-CLEM) system named HOPE-SIM by incorporating a 3D structured illumination fluorescence microscopy (SIM) system and an upgraded high-vacuum stage to achieve efficiently targeted cryo-FIB. With the 3D super resolution of cryo-SIM as well as our cryo-CLEM software, 3D-View, the correlation precision of targeting region of interest can reach to 150 nm enough for the subsequent cryo-lamella fabrication. We successfully utilized the HOPE-SIM system to prepare cryo-lamellae targeting mitochondria, centrosomes of HeLa cells and herpesvirus assembly compartment of infected BHK-21 cells, which suggests the high potency of the HOPE-SIM system for future *in situ* cryo-ET workflows.

## INTRODUCTION

Revealing three-dimensional (3D) cellular ultrastructures is an important step in understanding life. In recent years, cryo-electron microscopy (cryo-EM) has become increasingly improved and is becoming one of the major biophysical techniques used to study high-resolution 3D structures of macromolecular complexes ^1^. In addition, cryo-electron tomography (cryo-ET) has emerged as another powerful tool to visualize the macromolecular organization of unperturbed cellular landscapes with the potential to attain near-atomic resolution ^2,3^. To visualize the subcellular ultrastructure in vitrified cells by cryo-ET, focused ion beam milling under cryogenic conditions (cryo-FIB) has been used to prepare thin cryo-lamellae from vitrified cells, enabling many exciting biological observations to be made inside cells. The three-dimensional organization of cellular organelles, such as the cytoskeleton ^4-6^, endoplasmic reticulum (ER) ^7^, and 26S proteasome ^8^ have been successfully analyzed by cryo-ET.

There are several limitations that preclude the wider application of cryo-ET, including difficulties in locating and identifying features of interest ^9^. Cryo-correlative light and electron microscopy (cryo-CLEM) ^10-15^ has been proven to be an effective approach to overcoming this problem by utilizing fluorescent labeling to navigate toward the target for the subsequent cryo-FIB milling of vitrified cells ^11-14^ or cryo-ET imaging of frozen hydrated sections ^9,15^. Considering that the normal thickness of cells is far beyond the mean free path of 300 keV electrons, the fabrication of vitrified cells to a thickness of ∼200 nm is necessary before cryo-ET data collection. Therefore, the sequential experimental procedure from vitrification, cryo-fluorescence microscopy (cryo-FM), cryo-FIB and cryo-ET has become a routine workflow for many site-specific *in situ* structural studies (**Fig. 1**) ^9,16-20^. This workflow normally requires a stand-alone fluorescence microscope with a cryo-stage for cryo-fluorescence imaging ^12-14,21,22^, followed by cryo-scanning electron microscopy (cryo-SEM). A correlation alignment between the cryo-FM and cryo-SEM images is generated and used to guide cryo-FIB milling ^9,15,18,23^. In addition, specific fiducial markers imaged by both cryo-FM and cryo-FIB, such as fluorescent beads, are required for correlation alignment using specific 3D correlative software ^18,24^. There have been many hardware implementations reported to realize this workflow by developing cryogenic widefield fluorescence microscopy ^10,18,19^ or high-resolution cryo-Airyscan confocal microscopy (CACM) ^9^.

**Figure 1.**
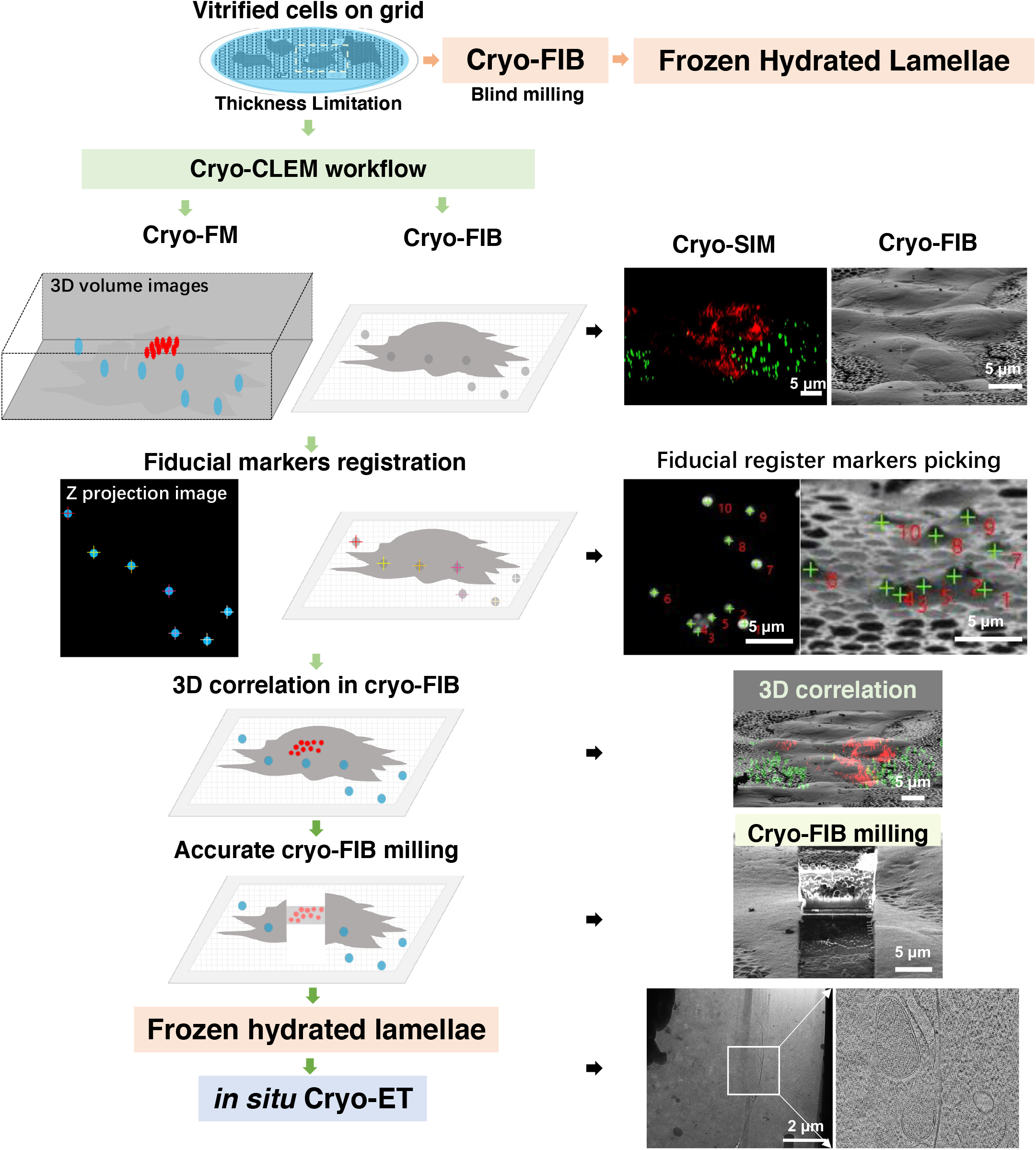
Workflow of cryo-CLEM with one example using cryo-SIM. Vitrified cells on EM grids are too thick to be imaged directly by cryo-EM. Blind milling by cryo-FIB lacks the precision to target the region of interest (ROI), which needs to be solved by cryo-CLEM. The vitrified cells were first imaged by cryo-FM and screened with good fluorescent signals. The fluorescent images of both microspheres (blue) and ROIs were captured. The fluorescent images are then correlated with cryo-FIB images by aligning the positions of microspheres (Fiducial marker registration). The precise correlation is then used to guide accurate cryo-FIB milling to prepare target cryo-lamellae of the ROI for the subsequent *in situ* cryo-ET study. An example using cryo-SIM of the HOPE-SIM system to demonstrate the whole workflow of cryo-CLEM is shown in the right panels. HeLa cells stained with MitoTracker Red CMXRos (red channel) were used for this demo. The fluorescent microspheres were imaged in the green channel and correlated with the cryo-FIB image, where the microspheres are marked with crosses. Accurate cryo-FIB milling was performed with the guidance of 3D correlation, and the subsequent cryo-ET reconstruction showed the location of the target mitochondria *in situ*.

The correlation accuracy and success rate between cryo-FM and cryo-FIB is limited by the resolution of cryo-FM and further attenuated by the factors of specimen deformation, devitrification, and ice contamination ^7,21^. We previously developed a unique <u>h</u>igh-vacuum <u>o</u>ptical <u>p</u>latform for cryo-CL<u>E</u>M (HOPE) ^22^, which utilized an integrative cryo-holder and a specific vacuum chamber to minimize ice contamination and reduce the risk of specimen deformation and devitrification during cryo-FM imaging and specimen transfer. However, the use of a widefield fluorescence microscope with a low numeric aperture (NA) objective lens and without the information of the z position of fluorescent targets in our HOPE system has limited the correlation accuracy above one micron, which needs to be efficiently improved to increase the success rate of site-specific cryo-FIB milling.

Structured illumination microscopy (SIM) introduces patterned illumination light on the imaging target, resulting in low-frequency interference fringes that carry structural information beyond the diffraction limit of the system, which doubles the resolution of conventional wide-field fluorescence microscopy and enables the visualization of detailed molecular processes in the whole cell ^4,25,26^. Compared to other superresolution FM techniques, such as photoactivated localization microscopy (PALM) ^27^, stochastic optical reconstruction microscopy (STORM) ^28^, stimulated emission depletion (STED) nanoscopy ^29^ and 4Pi microscopy ^30^, SIM uses relatively low illumination power (10–100 mW/cm^2^) and requires no special fluorophores or objective lenses to double the diffraction-limited resolution in three dimensions ^25,31-35^. The low illumination power minimizes the heating effect on the specimen, which is important for avoiding the risk of devitrification ^36,37^ for frozen hydrated specimens. Therefore, SIM has been recently employed within correlative microscopy with soft X-ray microscopy to visualize cells in vitreous ice ^32^.

In this work, starting from our previously developed HOPE system ^22^, we developed a new cryo-correlative light and electron microscopy (cryo-CLEM) system with the name HOPE-SIM to achieve efficiently targeted cryo-FIB. We upgraded the wide-field fluorescence microscope to a 3D-SIM system to increase the FM imaging resolution in the x/y dimensions and obtain additional information in the z dimension, which greatly improved the correlation accuracy to guide site-specific cryo-FIB fabrication. We also upgraded the high vacuum system to improve the vacuum of the chamber and further reduce the rate of ice growth, which allows a longer time of cryo-FM imaging before the specimen undergoes frosting. We also developed a new 3D correlative software, 3D-View, to perform the fiducial marker-based correlation between cryo-SIM and cryo-FIB images, which is used to navigate cryo-FIB milling accurately.

With our HOPE-SIM system, the frozen hydrated specimen can be transferred from cryo-SIM to cryo-FIB and then to cryo-ET without touching the grid directly, minimizing the risk of specimen deformation and contamination. The correlation precision of targeting region of interest can reach to 150 nm, which was further assessed by real experimental applications to target mitochondria, centrosomes of HeLa cells and herpesvirus assembly compartment of infected BHK-21 cells.

## RESULTS

### Design of the HOPE-SIM system

Our HOPE-SIM system (**Fig. 2a and Supplementary Fig. 1; Supplementary Video 1**) was upgraded from our previously developed HOPE system ^22^. First, a larger high vacuum chamber was designed and mounted onto a commercial epi-fluorescence microscope (e.g., model IX73, Olympus Corporation, Japan) with a high numerical aperture (NA) long working distance dry objective lens (e.g., Nikon CFI TU Plan Apo EPI 100×, NA/WD: 0.9/2 mm) mounted in the chamber (**Fig. 2a and Supplementary Fig. 1**). Placing the objective lens inside the vacuum chamber allows for the utilization of a high NA lens to increase the FM resolution. Similar to our previous design, a vacuum transfer system with a pre-pumping device is connected to the vacuum chamber via a corrugated tube and a manual gate valve, which allows a commercial cryo-holder to be transferred into the vacuum chamber and the cryo-specimen to be placed on top of the objective lens. The movement of the cryo-holder is driven by a three-axis electric motor stage that is fixed to the vacuum chamber. Different from our previous HOPE system, a new anti-contamination system (ACS) was designed, fixing a liquid nitrogen dewar in the vacuum chamber to cool a cryo-box that is placed on top of the objective lens (**Fig. 2a and Supplementary Fig. 1b**). This cryo-box with a working temperature lower than −175 degrees is used to cage the tip of the cryo-holder (**Supplementary Fig. 1c-e**) and protect the cryo-specimen from ice contamination. In addition, there are two ports designed on the left of the vacuum chamber, with one port connected to a turbo pump system (TPS) and another connected to a vacuum gauge (**Supplementary Fig. 1a-b**). A high-power TPS is used to achieve an improved vacuum better than 5×10^−5^ Pa under cryogenic conditions, which is important for reducing the ice growth rate during the period of cryo-FM imaging. We designed a touch screen device to monitor the vacuum levels of both the chamber and the gate valve, the pumping status, and the temperature of the cryo-box (**Supplementary Fig. 1a**). To monitor the system around the objective lens inside the chamber, we designed an observation glass window in front of the chamber (**Supplementary Fig. 1a, e**). To allow the light to pass through the vacuum chamber, similar to the original design of HOPE, there are two glass optical windows designed on the top and bottom of the objective lens (**Supplementary Fig. 1f, g**).

**Figure 2.**
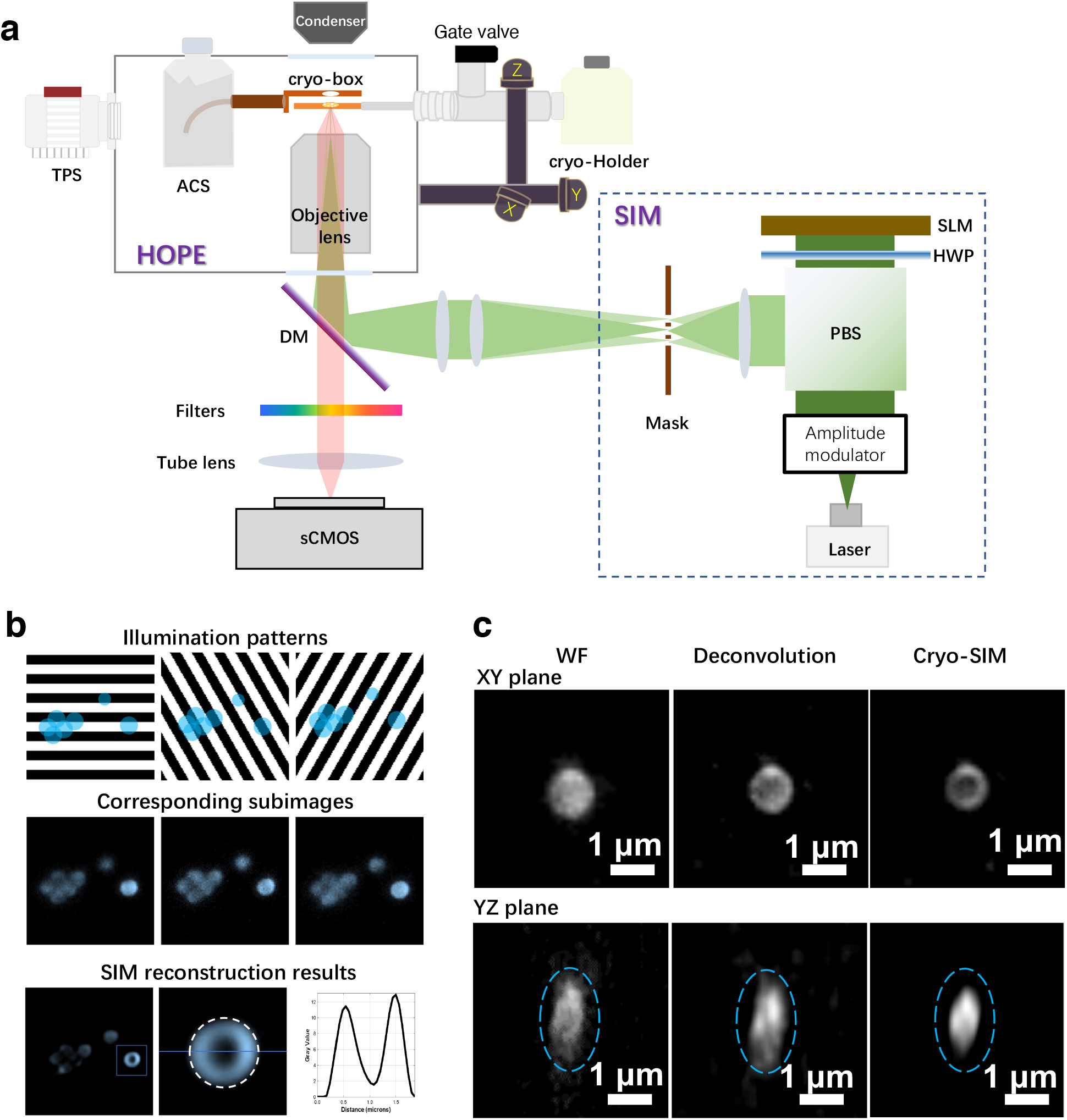
Design and principle of the HOPE-SIM system. (a) Schematic overview of the HOPE-SIM system design in its operational mode. Each part of the system is labeled and described in the text. A structured illumination beam is induced into the microscope via the back light inlet. TPS, turbo pump system; ACS, anti-contamination system; DM, dichroic mirror; SIM, structural illumination microscopy; SLM, spatial light modulator; HWP, half-wave plate; PBS, polarizing beam splitter. (b) Diagrammatic (top) and real (middle) corresponding subimages of the cryo-SIM illumination patterns (black and white) on the fluorescence microspheres (blue). SIM reconstruction results (bottom) show a hollow feature of a 1 μm microsphere. (c) Comparisons of fluorescent images of a 1 μm microsphere among different imaging modes: wide-field (WF), deconvolution and cryo-SIM. It is clear that cryo-SIM yields the image with the best resolution in both the XY and YZ planes.

To avoid touching the grid directly in the subsequent cryo-FIB and cryo-ET, we selected FEI AutoGrid for cryo-specimen transfer during the whole cryo-CLEM workflow. A multi-specimen single tilt cryo-holder (Model 910.6, Gatan, USA) was used to transfer the cryo-specimen into the vacuum chamber and maintain the cryogenic condition during cryo-FM imaging. To fit AutoGrid with this commercial cryo-holder, we used our previously reported holder tip ^38^ (**Supplementary Fig. 1c, d**) to hold AutoGrid mounted with our D-shaped finder grids ^39^ (**Supplementary Fig. 1c**). This special holder tip is important in protecting cryo-specimens from damage and providing enough mechanical stability for subsequent cryo-FM imaging.

In addition to upgrading the vacuum system and mechanics, we also upgraded the wide-field fluorescence microscope of our HOPE system to SIM (**Fig. 2 and Supplementary Fig. 2**), which utilizes a laser source with three channels (561 nm/488 nm/405 nm) and a spatial light modulator (SLM, QXGA-3DM, Forth Dimension Displays) to generate a structural illumination pattern. The resulting fluorescence signals are collected by the objective and recorded on a high-sensitivity sCMOS camera (Prime 95B, Photometrics). 3D-SIM raw data can be obtained to reconstruct a stack of fluorescence image series that will be used for the subsequent correlation and cryo-FIB navigation.

To control the whole system (laser, SLM, stage and camera) of HOPE-SIM and be compatible with the subsequent correlation, we designed a new LabVIEW (National Instrument, USA)-based software, HOPE-SIM View (**Supplementary Fig. 3**). HOPE-SIM View can be used to set up cryo-FM imaging parameters (e.g., the exposure time and focusing zone), map the area of the whole grid, register the list of targets of interest (**Supplementary Fig. 3a**), change the laser channel and power intensity (**Supplementary Fig. 3b**), control the stage movement (**Supplementary Fig. 3c**), and collect multichannel 3D-SIM raw data.

### HOPE-SIM-based cryo-CLEM workflow

Using our HOPE-SIM system, we can incorporate cryo-SIM imaging into the current standard protocol of cryo-CLEM (**Fig. 1**). In detail, the D-shaped finder grid is used to grow cells to a suitable density, and the cells are then either transfected to express target florescent tagged molecules or stained with fluorescent dyes. Before plunge freezing, a proper amount of diluted fluorescent microspheres with a recommended size of 1 micron in diameter are added into the culture on the grid. The vitrified grid is then assembled into AutoGrid and mounted into our specially designed tip that is fitted into the multispecimen cryo-holder for the subsequent cryo-SIM experiment. Notably, the side of the grid with the growing cells should face down to adapt to the working distance between the cells and objective lens.

Before cryo-SIM imaging, the vacuum chamber of the HOPE-SIM system must be pumped to a high vacuum of ∼5×10^−5^ Pa, and the ACS needs to be cooled to less than −170 degrees by liquid nitrogen. The cryo-multiholder is then inserted into the HOPE-SIM system by the vacuum transfer device. After ∼5 minutes of waiting time to recover the vacuum of the chamber and another ∼5 minutes for the temperature of the holder tip to stabilize, the shutter of the multiholder is open and ready for sample screening and cryo-SIM imaging. The frozen grid is first screened in transmission wide-field imaging mode, and the concentration of cells, positions of cells in the square, integrity of the supporting film, ice contamination and thickness of the specimen are checked. The regions with cells showing a good fluorescent signal at the right position in the square, an unbroken supporting film, an appropriate ice thickness, and well-dispersed fluorescent beads are selected and registered as ROIs (regions of interest) in HOPE-SIM View for the subsequent 3D cryo-SIM data collection (**Supplementary Fig. 3**), which covers a whole square of a 200-mesh grid and includes channels of both fluorescent beads and target fluorescent molecules.

After cryo-SIM imaging, the frozen grid assembled with AutoGrid is transferred into a dual beam cryo-SEM (e.g., FEI Helios NanoLab 600i) via our previously developed protocol and cryo-shutter ^39^. We developed LabVIEW (National Instrument, USA)-based software, 3D-View (**Supplementary Fig. 4**), for image correlation between different microscopes (**Supplementary Video 2**). First, the whole grid is screened by cryo-FIB imaging in low magnification, and the positions of the ROIs can be found via the index of the Finder. The ROIs with cells having good shape and suitable density are selected for subsequent cryo-FIB imaging with a proper magnification of 2,500X (**Supplementary Fig. 4a-b**). Then, the positions of fluorescent beads from both the cryo-SIM and corresponding cryo-FIB images are selected for precise correlation using 3D-View (**Supplementary Fig. 4b**) (the detailed algorithm will be described in the next section). After image correlation, the optimized Euler angle and minimized standard deviation (SD) can be obtained and displayed (**Supplementary Fig. 4c-e**). Then, the fluorescence image of cryo-SIM can be merged with the cryo-FIB image (**Supplementary Fig. 4f**) to navigate to the target for the subsequent fabrication of cryo-lamella with a thickness of ∼200 nm.

Using AutoGrid, the prepared cryo-lamella is then ready for the cryo-ET experiment using a 300 kV cryo-electron microscope (e.g., FEI Titan Krios). The cryo-SIM image is also useful to help locate the ROI for cryo-ET data collection.

### Development of 3D-View for image correlation

In 3D-View, we developed an algorithm to correlate the 3D cryo-SIM image and 2D cryo-FIB image of fluorescent microspheres (**Supplementary Video 2**). The transformation (rotation and translation in three dimensions) between cryo-SIM and cryo-FIB images is computed based on the coordinates of the corresponding fluorescent microspheres in two images ^18^. The coordinates of geometric centers are important for the final correlative accuracy. The z coordinates of fluorescent microspheres were determined from the cross-section of cryo-SIM 3D volume data of the fluorescent microspheres (**Fig. 2c & Supplementary Fig. 4d**).

We defined the coordinate vector of each fluorescent microsphere a 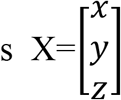; this provided two datasets of vectors, *X*_*SIMi*_ (*i* = 1, …, *N*) and *X*_*FIBi*_ (*i* = 1, …, *N*), for the selected corresponding fluorescent microspheres from the cryo-SIM and cryo-FIB images, respectively. The centr oid of each dataset can be calculated as follows:

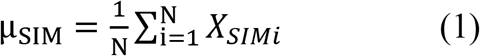

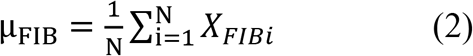

In this study, we defined the rotation operation R(θ) of three Euler angles (θ_*x*_, θ_*y*_, θ_*z*_) as follows:

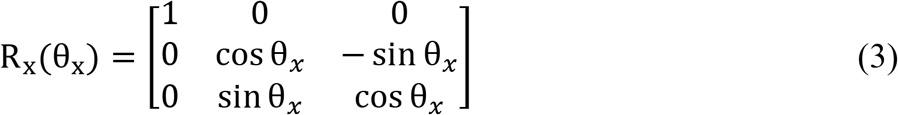

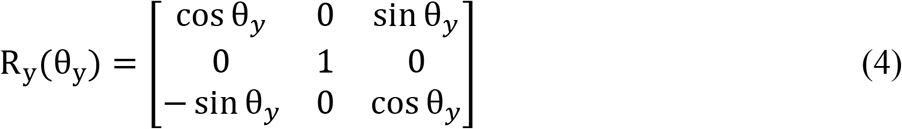

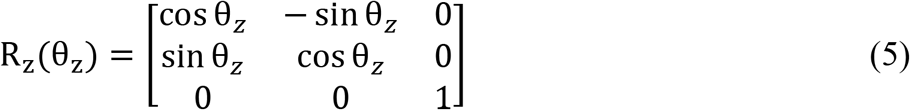

The 3D coordinates of the fluorescent microspheres in the cryo-SIM volume can be translated into 2D coordinates on the coordinate plane of the cryo-FIB image as follows:

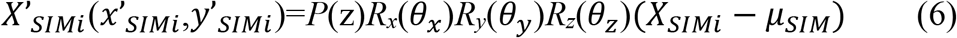

where P(z) is the projection along the z direction and the Z-Y-X rotation transformation system is used.

Additionally, the coordinates of the fluorescent microspheres in the cryo-FIB image are translated as follows:

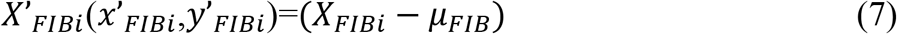

According to Equations (6) and (7), the deviation of the correlation between the cryo-SIM and cryo-FIB datasets can be defined as follows:

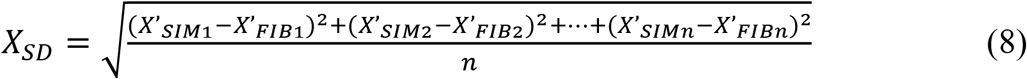

Using the least square methods, the optimized Euler angle (θ_*x*_, θ_*y*_, θ_*z*_) can be solved to minimize the correlation deviation *X*_*SD*_. Finally, the cryo-SIM 3D volume data can be projected and merged with the 2D cryo-FIB image using Equation (6). The target positions defined in the cryo-SIM image can be located precisely on the cryo-FIB image.

We implemented the above mathematical algorithm into our 3D-View software to correlate multichannel 3D cryo-SIM images with 2D cryo-FIB images in a semi-automatic way.

### Precision of targeted cryo-FIB

Using the protocol of the open-source software SIM-Check ^40^, we measured the resolution and checked the stability/quality of our cryo-SIM system (**Supplementary Fig. 2e-i**). As a result, an ∼200 nm lateral resolution and an ∼500 nm z-direction resolution were achieved (**Supplementary Fig. 2e, f**), which are better than the wide-field image as well as the deconvolution image (**Fig. 2c**). The enhanced resolution and signal-to-noise ratio improved the accuracy of the determined z-coordinates of the fluorescent microspheres, which is important for the subsequent correlation accuracy between the 3D cryo-SIM and 2D cryo-FIB images.

We then measured how precise cryo-FIB milling could be to the fluorescent target by using another color of fluorescent microsphere as a test target (**Fig. 3**). First, two differently sized and colored fluorescent microspheres (green/1 μm, red/0.5 μm) were spread onto a GraFuture RGO TEM Grid (GraFuture™-RGO001A, Shuimu BioSciences Ltd., CN) and then dried. We used the channel of green fluorescent microspheres to correlate the 3D cryo-SIM and 2D cryo-FIB images and then calculated the standard deviation of the coordinates of the red fluorescent microspheres according to Equation (8), resulting in an ∼150 nm correlation precision (**Fig. 3a-c**). Second, to mimic the real experimental procedure of target cryo-FIB milling, we vitrified the solution of green fluorescent microspheres directly on the EM grid, utilized the microspheres on the ice surface as markers to correlate the 3D cryo-SIM and 2D cryo-FIB images, and then took the microspheres embedded in the ice as the target for cryo-FIB milling (**Fig. 3d-f**). After target cryo-FIB milling, cryo-lamella with a thickness of ∼200 nm was then transferred for cryo-EM imaging. We then projected the original 3D cryo-SIM image onto the cryo-EM micrograph and found that the target microsphere was accurately localized in the center of the micrograph, which shows that cryo-FIB milling navigated by correlation with the 3D cryo-SIM image can reach the target with a precision better than 200 nm.

**Figure 3.**
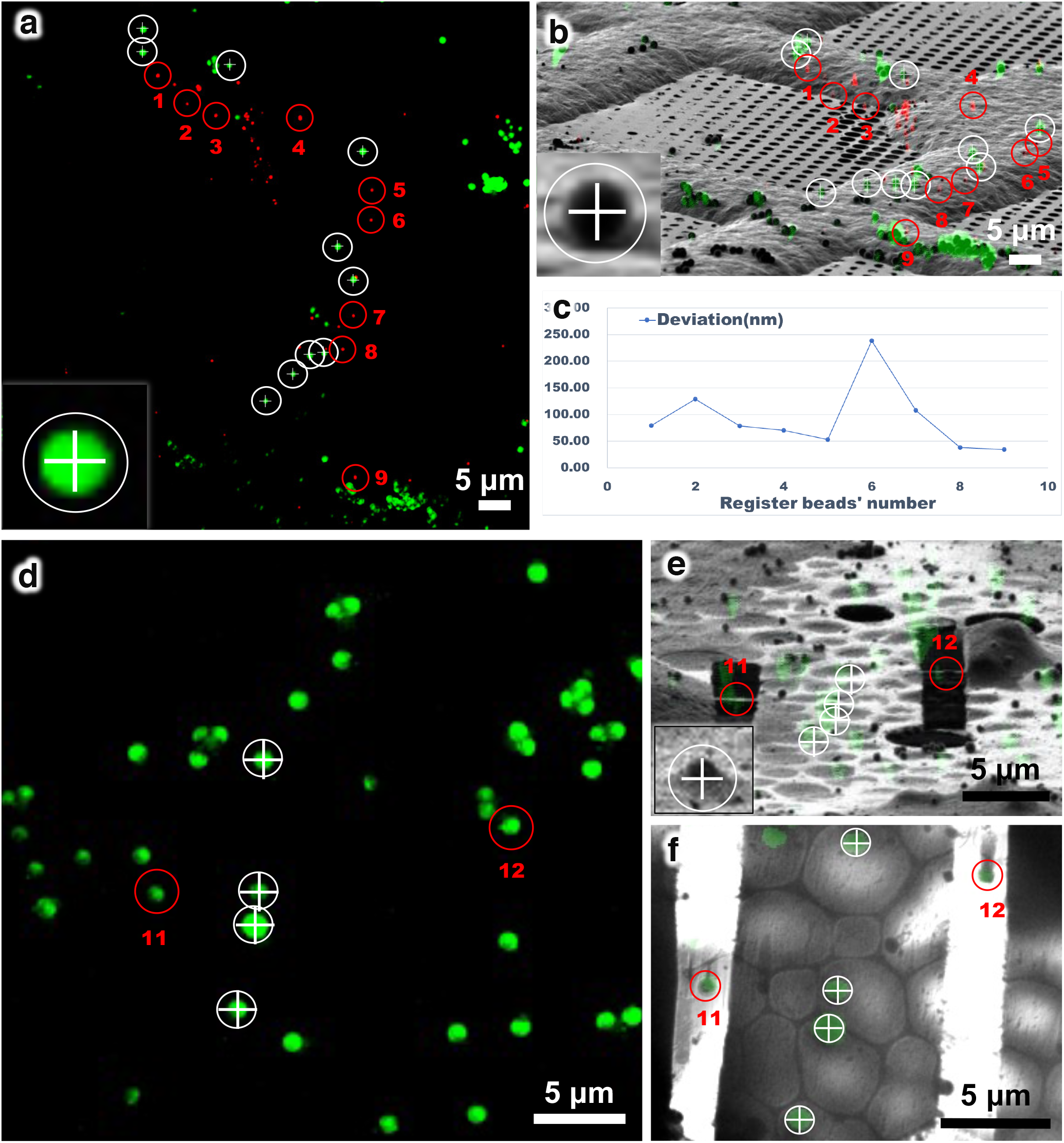
Correlation accuracy of the HOPE-SIM system. (a) HOPE-SIM images of dried 1 μm green and 500 nm red fluorescent microspheres spread on a GraFuture RGO grid. The green microspheres tagged by white circles and crosses are used as the fiducial register markers. The red microspheres labeled from 1 to 9 are used to measure the correlation accuracy with the FIB image. (b) 3D correlation with the FIB image. Microspheres corresponding to the green microspheres in (a) are tagged by white circles and crosses and used to perform correlation with the fluorescent image. Microspheres tagged with red circles and labeled from 1 to 9 correspond respectively to the red microspheres in (a). (c) Position deviations of nine corresponding red microspheres between (a) and (b) after the correlation is plotted. (d) HOPE-SIM image of a vitrified solution of fluorescent microspheres dropped directly on the EM grid. The microspheres tagged with white circles and crosses are used to perform image correlation. Those tagged with red circles and labeled as 11 and 12 are used to verify the accuracy of the subsequent site-specific cryo-FIB milling. (e) 3D correlation with cryo-FIB image for site-specific milling. The corresponding microspheres in (d) are found and marked with white circles and crosses. These microspheres are now embedded in ice and appear as bumps in the cryo-FIB image. Their centers are then determined by placing the top end of the cross on the top of the bump, and the length of the crossbar is set to the diameter of the microsphere. Then, an accurate 3D correlation between 3D cryo-SIM and 2D cryo-FIB images can be performed. The target positions labeled 11 and 12 are subjected to cryo-FIB milling to make two cryo-lamellae with thicknesses of ∼200 nm. (f) Cryo-EM micrograph (450X) of the cryo-lamellae is merged with the fluorescence image in (d). The target microspheres 11 and 12 in (d) can be clearly visualized in the micrograph and kept in the cryo-lamellae.

Notably, we found that the distribution of the fluorescent microspheres in the y direction (perpendicular to the FIB scanning direction) and the precision of determination of the z-coordinates of the microspheres are key factors affecting the accuracy of the correlation (see also **Supplementary Note 1**).

### Application of *in situ* ultrastructural study

We applied our HOPE-SIM-based cryo-CLEM workflow to study real biological systems. The first test was to target the mitochondria of HeLa cells that were stained with MitoTracker Red CMXRos (Invitrogen M7512, Thermo Fisher Scientific Corp., OR.). With the improved resolution of cryo-SIM, the mitochondria can be well resolved and separated in the fluorescent image, and the selected mitochondrion can be precisely targeted by 3D-View for cryo-FIB milling, which is subsequently captured by cryo-ET (**Fig. 1**).

In addition to targeting mitochondria, we tried a second test with more biological implications to visualize virus assembly *in situ* from infected cells. Herpesviridae is a large family that consists of alpha, beta, and gamma herpesviruses. Murine gamma herpesvirus 68 (MHV-68), which is similar to human gamma herpesviruses, can establish a productive infection of fibroblast or epithelial cell lines derived from mammalian species and therefore is a model system for the study of pathogenic mechanisms associated with human gamma herpesvirus infection. Herpesvirus maturation includes four stages: capsid formation in the nuclei, primary envelopment and passing through the nuclear membrane, tegumentation and secondary envelopment at the cytoplasm, and egressing from the cell ^41^. Using conventional transmission electron microscopy of plastic-embedded sections, previous studies observed ‘tegument deposits’ in the cytoplasm of MHV-68-infected BHK-21 cells, where capsids accumulate to acquire a series of tegument proteins ^42^.

Here, we labeled the capsid protein ORF65 of MHV-68 with the fluorescent tag mCherry and utilized our HOPE-SIM-based cryo-CLEM workflow to investigate the native *in situ* structure of the ‘tegument deposit’ in ORF65-mCherry MHV-68-infected BHK-21 cells (**Fig. 4**). We used 1 μm blue microspheres as fiducial markers to correlate 3D cryo-SIM and 2D cryo-FIB images in 3D-View and identified the tegument deposit location in the cytoplasm (**Fig. 4a**). Then, we performed site-specific cryo-FIB milling and prepared cryo-lamella with a thickness of 200 nm. We took a low magnification (4,300X) cryo-EM micrograph of the cryo-lamella and merged it with the corresponding cryo-SIM image (**Fig. 4b**), confirming that the localizations of viral particles precisely correlated with the fluorescent signal. We selected one region to take higher magnified cryo-EM micrographs and found different types of viral particles that were under tegumentation (**Fig. 4c**).

**Figure 4.**
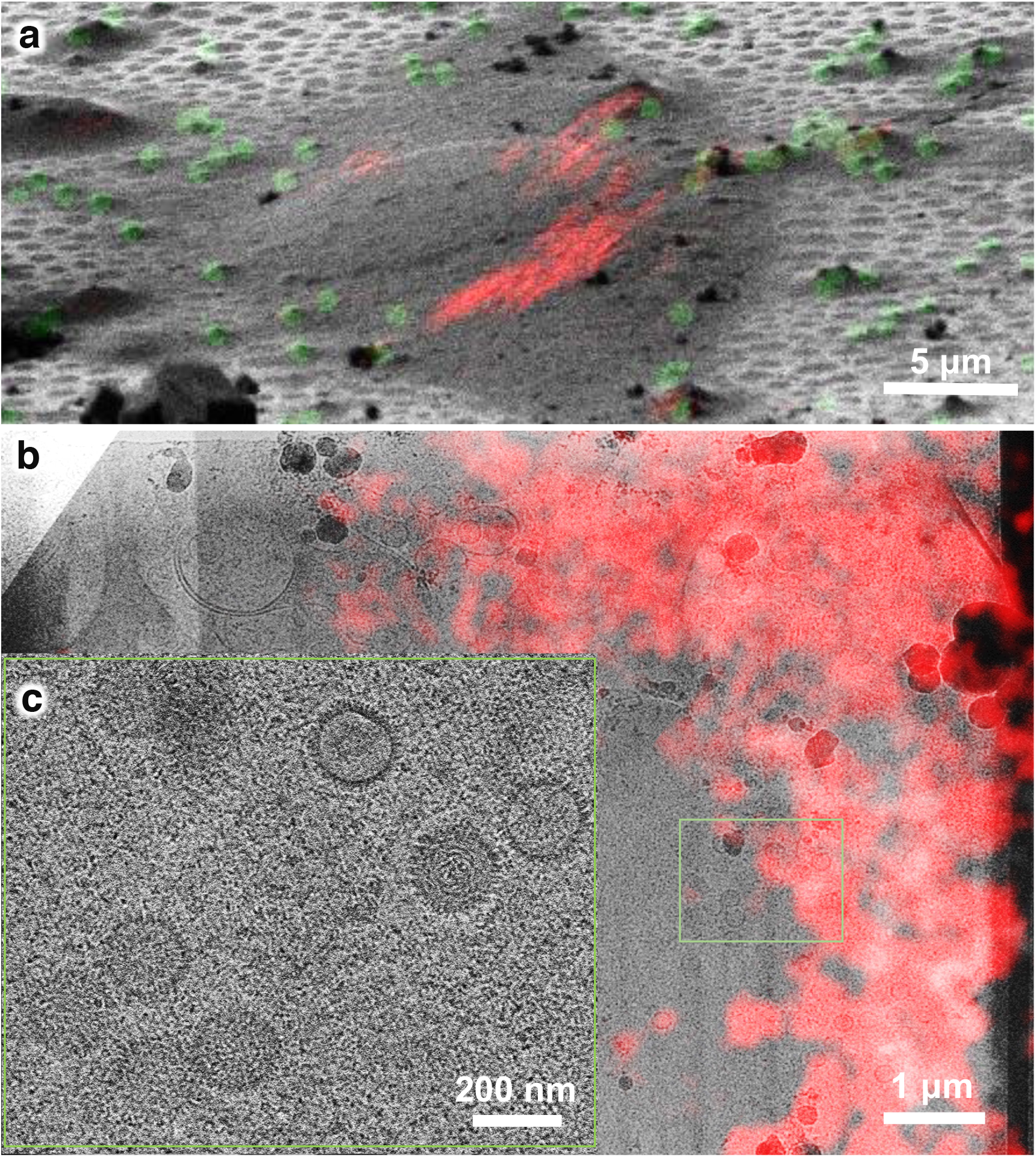
Using the HOPE-SIM-based cryo-CLEM workflow to capture the MHV-68 viral particles in host cells. (a)3D correlative image indicating the target region of the MHV-68 virus assembly compartment inside the host cell. Green, fluorescent microspheres. Red, MHV-68 viral particles. (b)Cryo-EM micrograph (4,300X) of the cryo-lamella is merged with the cryo-SIM image to show the location of the MHV-68 viral particles. The target region marked by the green rectangle is subjected to cryo-ET data collection and reconstruction. (c)One slice of the tomogram (33,000X) of the target region in (b), showing different types of viral particles that were under tegumentation.

For the third test, we tried a more challenging target, namely the human centriole in HeLa cells. Human centrioles are closely related to tumorigenesis and multiple hereditary diseases ^43,44^. There are only one or two pairs of centrioles in each mammalian cell, which greatly limits their *in situ* structural study without an efficient cryo-CLEM workflow. Here, we used HeLa cells expressing the mCherry-tagged PACT (pericentrin-AKAP450 centrosomal targeting) domain ^45^, which is one of the centrosome components, to target and visualize the human centriole by cryo-ET (**Fig. 5**). Again, we used 1 μm blue microspheres as fiducial markers to correlate the 3D cryo-SIM and 2D cryo-FIB images in 3D-View and identify the location of the centrosomes, appearing as two fluorescence dots in the cryo-SIM images (**Fig. 5a**). Then, due to the precise correlation, we performed site-specific cryo-FIB milling to prepare cryo-lamella containing centrosomes with a thickness of 200 nm (**Fig. 5b-d**), which was subsequently verified by cryo-EM (**Fig. 5e**). Further reconstruction by cryo-ET revealed the 3D *in situ* structure of a pair of perpendicular centrioles (**Figs. 5f-j and Supplementary Videos 3-5**). Notably, according to our current statistics, we managed to fabricate 19 thick cryo-lamellae from 19 vitrified cells and put back for cryo-SIM verification and we found 16 cryo-lamellae retained the fluorescence signal of target centrosomes (**Supplementary Fig. 5, Supplementary Data. 1 and Supplementary Table 1**), suggesting a high success targeting rate.

**Figure 5.**
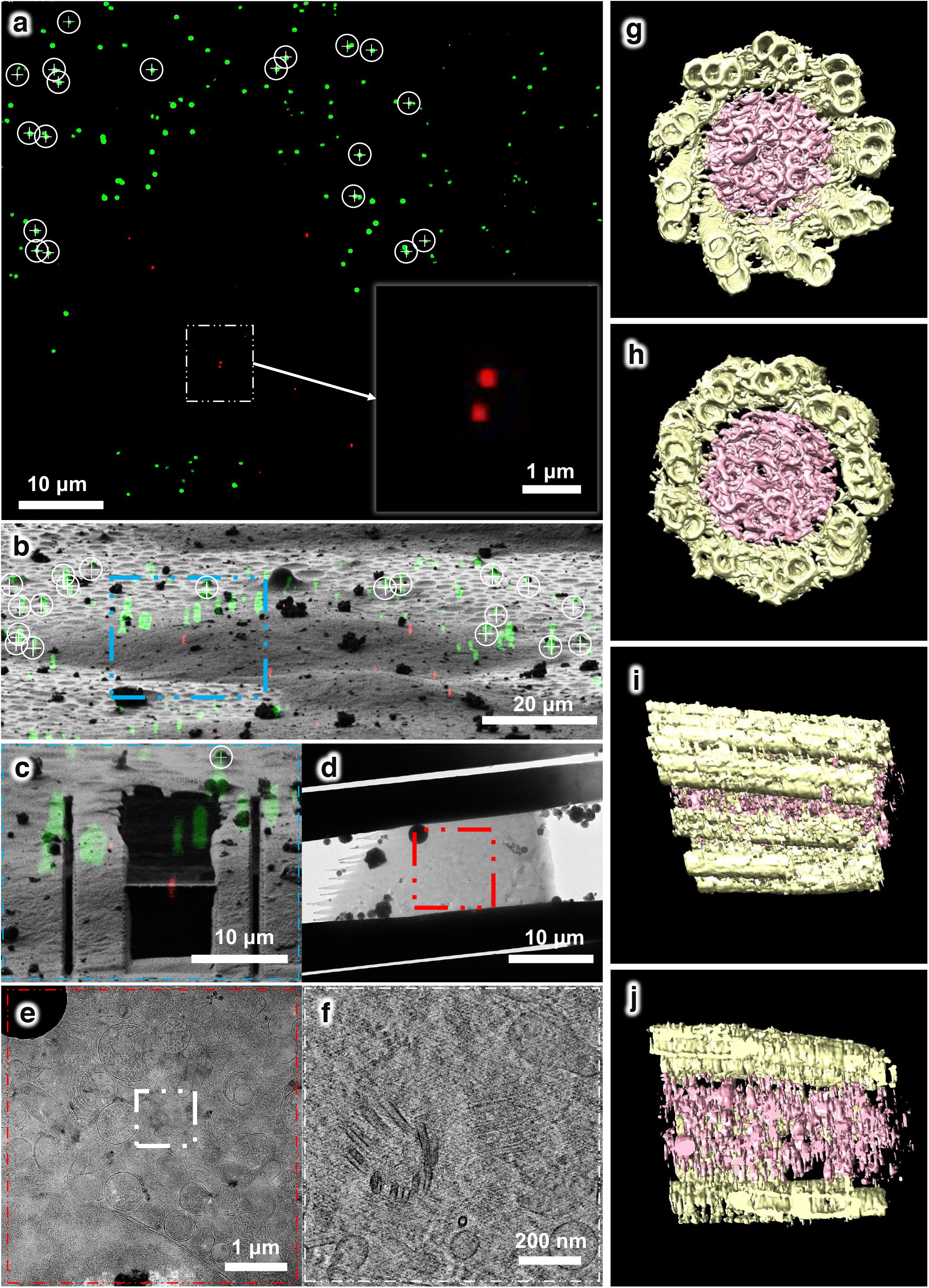
Using the HOPE-SIM-based cryo-CLEM workflow to capture the centrosomes in HeLa cells. (a)Cryo-SIM images indicate the well spread microspheres (green) and the fluorescence-labeled centrosomes (red). The ROI containing a pair of centrosomes is marked with the white square. The microspheres used as fiducial markers for image correlation are marked by white circles, with their centers indicated by white crosses. (b)3D correlative image indicating the position of the fluorescence-labeled centrosomes (red) inside the HeLa cell. The ROI in (a) is marked by an blue square. Green, fluorescent microspheres. The corresponding microspheres in (a) are marked by white circles, with their centers indicated by white crosses. (b)3D correlative image in the ROI after site-specific cryo-FIB milling. The fluorescent signal of the centrosomes (red) is located on the cryo-lamella. (c)Cryo-EM micrograph (4,300X) of the cryo-lamella in (c), which contains the target centrosomes shown in the red square. (d)Cryo-EM micrograph (8,700X) of the cryo-lamella in the region of the red square in (d). The target region marked by the white square is subjected to cryo-ET data collection and reconstruction. (e)One slice of the tomogram (53,000X) of the target region in the white square shown in (e). The target centrioles are shown with one in the top view and another in the side view. (g-j) 3D in situ structure of the centriole in HeLa cells with different views,(g)top, (h) bottom, (i) side and (j) cross. The structure is segmented using EMAN2 ^53^ and rendered in Chimera ^54^. The triplet tubules are shown in yellow, and the internal scaffold structures are shown in pink. This structure was derived from another tomogram (26,000X) of cryo-lamella with a thickness of ∼300 nm.

## DISCUSSION

*In situ* structural study of macromolecules in their native cellular context by cryo-ET has become a new frontier of structural biology. Subsequently, the demand for improving the overall workflow, from specimen preparation and data collection to image analysis, has increased. Additionally, the current protocol of image correlation among cryo-FM, cryo-FIB and cryo-ET needs to be further improved with better efficiency and accuracy. The commonly used wide-field cryo-FM ^10,18,19^ or recently developed cryo-Airyscan confocal microscopy (CACM) ^9^ have limited resolution and lack 3D resolution, which hampers the correlation accuracy with the subsequent 2D image from cryo-FIB. Superresolution fluorescence microscopy under cryogenic conditions based on the photon-activated localization principle has been developed with a nanometer-scale resolution ^37,46^; however, the high illumination power that is required increases the high risk of devitrification. Compared to conventional wide-field fluorescence microscopy, SIM enables a twofold improvement in the diffraction-limited resolution in three dimensions ^25,31-35^ with a relatively low illumination power (10–100 mW/cm^2^). The wide-field nature of SIM makes it light-efficient and decouples the acquisition speed from the size of the lateral field of view, meaning that high frame rates over large volumes are possible ^34^. Furthermore, SIM can be combined with a superresolution imaging algorithm ^35^ or a progressive deep-learning superresolution strategy ^47,48^ to achieve a higher resolution. Therefore, cryo-SIM provides a promising solution to improve the accuracy of the current cryo-CLEM workflow.

In the present study, we developed a HOPE-SIM-based cryo-CLEM workflow by incorporating the SIM module into our previously developed HOPE system to enable high-resolution 3D cryo-FM, upgrading the original vacuum system to further eliminate ice contamination, and writing new HOPE-SIM View/3D-View software to control the hardware and perform image correlation semi-automatically. We demonstrated that our HOPE-SIM system can achieve a resolution of ∼200 nm in the lateral direction and ∼500 nm in the z direction. These results are better than the conventional wide-field cryo-FM and deconvolution modes. We verified the precision of the correlation between cryo-SIM and cryo-FIB as ∼150 nm, which is good enough to perform accurate site-specific cryo-FIB fabrication of cryo-lamella with a thickness of ∼200 nm. We applied our HOPE-SIM-based cryo-CLEM workflow to successfully target MitoTracker-stained mitochondria in HeLa cells, visualize mCherry-tagged MHV-68 virions under tegumentation in infected BHK-21 cells, and obtain a tomogram of the human centrioles in HeLa cells. The high success rate of targeting the human centrioles suggests the robustness and accuracy of our cryo-CLEM workflow.

Compared to the published and commercially available cryo-CLEM system, our HOPE-SIM-based cryo-CLEM system offers numerous advantages, including the use of a high-vacuum chamber with an embedded high NA objective lens kept at room temperature and an upgraded anticontamination system to allow long-time high-resolution cryo-FM imaging without the concern of the lens becoming frozen/damaged and of ice growth/contamination; with the specially designed holder tip to accommodate the AutoGrid and commercial cryo-multiholder, the cryo-specimen can be efficiently transferred from cryo-FM to cryo-FIB, and then to cryo-ET without touching the grid directly, which minimizes the risk of specimen damage and ice contamination during specimen transfer. With cryo-SIM imaging and the 3D correlation algorithm, we could achieve high-resolution fluorescence microscopy, especially with a better z-direction resolution, and then realize a precise 3D correlation with cryo-FIB 2D images, resulting in accurate site-specific cryo-FIB milling.

Overall, our HOPE-SIM system-based cryo-CLEM workflow provides an efficient solution to achieve accurate target cryo-FIB fabrication of cryo-lamella ready for site-specific cryo-ET study, which will have wide applicability during future *in situ* structural studies. In addition, our HOPE-SIM system is versatile and can be adapted for various fluorescence and electron microscopes by utilizing proper stages, cryo-holders, and cartridges. In the future, we will combine our HOPE-SIM system with the progressive deep-learning superresolution strategy to achieve a higher resolution and update the 3D-View software to realize fully automatic image correlation, which will make site-specific cryo-FIB milling more efficient and accurate.

## METHODS

### Cell culture and vitrification

Indexed ultraviolet sterilized grids with one quarter trimmed (D-grids, T11012SSF, TIANLD, China) were used for HeLa and BHK-21 cell culturing.

HeLa cells labeled by MitoTracker or expressing mCherry fused to the PACT (pericentrin-AKAP450 centrosomal targeting) domain ^45^ were seeded onto ultraviolet sterilized D-grids and cultured in complete DMEM supplemented with 10% fetal bovine serum, 1% penicillin and streptomycin. After 48 h of culture, the grids were subjected to subsequent plunge freezing.

BHK-21 cells were cultured in complete Dulbecco’s modified Eagle’s medium (DMEM) supplemented with 10% fetal bovine serum, 1% penicillin and streptomycin. Culture media was routinely changed every 2 days by replacement of half the old medium with new medium. The ultraviolet-sterilized D-grids were placed into a 12-well plate culture dish before cell seeding (carbon film side upward). Then, BHK-21 cells were seeded. After cells attached to the D-grids, they were infected with ORF65-mCherry MHV-68 virus (constructed by Xing Jia in Dr. Hongyu Deng’s laboratory) at an MOI of 100 for 1 hr. Then, the inoculum was removed and replaced with fresh DMEM plus 10% fetal bovine serum. After 24 h of infection, the D-grids were subjected to plunge freezing.

Specifically, 1.5 μL of blue fluorescent microspheres with a diameter of 1 μm (430/465 nm, F13080, Thermo Fisher Scientific Corp., OR.), used as the markers for correlation between the cryo-SIM 3D and cryo-FIB 2D images, were diluted with PBS in a 1:400 ratio and added onto the D-grid with the side where the cells were growing before cryo-vitrification.

### Optical path of the HOPE-SIM system

A schematic diagram of the optical path of the HOPE-SIM system is presented in **Fig. 2a**, and a picture of the real optical setup is shown in **Supplementary Fig. 2a**. We set up a laser combiner of three lasers with wavelengths of 405 nm (50 mW, OBIS 405LX, Coherent), 488 nm (200 mW, SAPPHIRE 488-200 CW, Coherent), and 561 nm (200 mW, SAPPHIRE 561-200 CW, Coherent). The laser beam emitted from the laser combiner passes through an acousto-optic tunable filter (Amplitude modulator, AOTF, AOTFnC-400.650-TN & MPDS4C-B66-22-74.158-RS controller, Quanta Tech) and then expands to a diameter of 25 mm. Then, the beam passes through a phase-and-angle modulator consisting of a polarizing beam splitter cube (PBS, CCM1-PBS251, Thorlabs), an achromatic half-wave plate (HWP, AHWP10 M-600, Thorlabs), and a ferroelectric spatial light modulator (SLM, QXGA-3DM, Forth Dimension Displays) to generate patterned illumination with three alternative orientations (**Supplementary Fig. 2b**). These three patterned beams pass through another lens and are modulated at the diffraction plane using a customized mask consisting of a lightproof disc with a seven-hole aperture. Only diffractions at 0 and ±1 orders are selected to pass for each patterned beam. All three patterned beams are eventually focused at the back focal plane of the objective lens (**Supplementary Fig. 2c**). After passage through the objective lens, interference patterns (Moiré fringe) are generated with five phases (0, π/5, 2π/5, 3π/5, 4π/5) and three rotation angles (0°, 60°, and 120°). The period, orientation, and relative phase of this excitation pattern can be finely tuned by adjusting the SLM setup. For each orientation and phase of the excitation pattern, the resulting fluorescence signals are collected by the objective lens and recorded on a high-sensitivity sCMOS camera (Prime 95B, Photometrics). All lenses used in the system are chosen as achromatic doublets. Optical apertures and masks are used to improve the illumination quality.

### 3D-SIM data collection and image reconstruction

Raw data serials of 3D-SIM using a dry objective lens (NA 0.9) were collected with an increment of 250 nm along the z-axis. For each z-slice, 15 images were acquired for five phases and three orientations to satisfy Nyquist-Shannon sampling. Each raw image has a pixel size of 120 nm and a field of view (FOV) of 1,200 × 1,200 pixels. Then, the final raw image stack has a voxel size of 120 × 120 × 250 nm. The exposure time for each raw image depends on the overall fluorophore intensity, and normally, the entire dataset was collected within an average time of 5-10 min to capture a field of view with the required z-depth of ∼20 μm SIM image reconstructions were performed using SoftWoRx 6.1.1 (GE Healthcare) with the following settings: channel-specific optical transfer functions (OTFs), a Wiener filter constant of 0.010, a background with negative intensities discarded, and drift correction with respect to the first angle. OTFs were generated from point spread functions (PSFs) by SoftWoRx 6.1.1, and PSFs of the HOPE-SIM system at cryogenic temperatures were measured by using 200 nm fluorescent microspheres (405 nm/488 nm/560 nm, Thermo Fisher Scientific Corp., OR.). For each z-slice, the final reconstructed image contains 2,400 × 2,400 pixels with a pixel size of 60 nm. The open-source software SIMcheck ^40^ was used to verify the performance of the SIM optical system, tune parameters and recognize artifacts.

### 3D correlative registration and cryo-FIB milling

The clearly visible fluorescent microspheres around the target cell were selected manually from the cryo-SIM 3D and cryo-FIB images. The distribution of the selected fluorescent microspheres should be on both sides of the cell, especially in the y direction, for better correlation accuracy. The centers of the microspheres in the XY plane were determined first. To further optimize the correlation between two correlative images, the Z height of fluorescent microspheres could be manually adjusted according to the shapes of the fluorescence microspheres (shown in **Supplementary Video. 2**). For the cryo-FIB image, the center of the microsphere was difficult to recognize because the microsphere was normally embedded in the ice and appeared as a bump. The position of the top of the bump represents the position of the microsphere edge. Therefore, with this knowledge, the center of the microsphere in the cryo-FIB image could be determined by finding the position that was a distance of half the diameter of the microsphere from the top of the bump along the y direction (FIB milling direction). After the first round of correlation, some microspheres with large deviations were removed due to the significant errors when determining their centers. The correlation parameters were then updated by optimizing the z-axis position of each microsphere. All these image correlation operations were performed in 3D-View (**Supplementary Fig. 4**).

Cryo-lamellae were prepared by cryo-FIB milling using the FEI Helios NanoLab 600i or Aquilos-2 FIB-SEM microscope (Thermo Fisher Scientific Corp., OR.). Target cells with clear fluorescent signals were selected according to the above correlation map. The grid was then coated with an organometallic platinum layer for 5 s at a 5 mm work distance. Cryo-lamellae with ∼200 nm thickness were obtained by milling gradually using a 4-step Ga^+^ beam current of 0.43 nA, 0.23 nA, 80 pA and 40 pA at a stage angle of 13– 22°.

### Cryo-electron tomography data collection and reconstruction

The cryo-lamellae were loaded into an FEI Titan Krios G3i (Thermo Fisher Scientific Corp., OR.) that was equipped with a Gatan GIF K2 4k × 4k camera (Gatan, Inc., Warrendale, PA) and operated at 300 kV in low-dose mode. Tilt series were collected bidirectionally with SerialEM software ^49^ using a tilt range of −60° to 40° in 2° intervals, nominal magnification of 33,000× or 53,000×, defocus of −4 ∼ ™6 μm and total dose of ∼150 e/Å^2^. The tilt series were aligned and reconstructed with a weighted back-projection algorithm using IMOD ^50^ or reconstructed using ICON ^51,52^.

### Segmentation and visualization

Image segmentation of the centriole tomogram was performed using EMAN2 ^53^, and 3D rendering was performed using Chimera ^54^.

## Data availability

The data that support the findings of this study are available from the corresponding author upon request.

## Code availability

The LabVIEW program HOPE-SIM View for controlling the device is hardware dependent, and the 3D-View program for image correlation is hardware-independent. These two programs are available at GitHub with the link, https://github.com/hilbertsun/HOPE-SIM.

## Acknowledgments

All cryo-CLEM, FIB and ET work was performed in the Center for Biological Imaging (CBI, http://cbi.ibp.ac.cn), the Institute of Biophysics (IBP), and the Chinese Academy of Sciences (CAS). We are grateful to Dr. Xiaojun Huang, Boling Zhu, Xujing Li, Lihong Chen for their help with cryo-EM data collection and Mr. Zeyang Li and Ms. Lulu Qin for their help with cryo-FIB fabrication. We would also like to thank Dr. Fulin Wang and Prof. Jianguo Chen from Peking University for their kindness in providing HeLa cells expressing mCherry-PACT, Prof. Wei Ji and Prof. Dong Li from IBP for their kind help with light microscopy, and Ms. Ziyan Wang and Dr. Yun Zhu from IBP for their kind help with cryo-ET data processing.

This work was equally supported by grants from the Ministry of Science and Technology of China (2017YFA0504700 to GJ), the Strategic Priority Research Program of Chinese Academy of Sciences (XDB37040102 to FS) and the National Natural Science Foundation of China (31830020 to FS). In addition, this work was supported by the Technological Innovation Program of the Chinese Academy of Sciences (29Y8CZ021001 to GJ), the CAS Key Technical Support Personnel Project (29Y9CQ041 to GJ), and the National Natural Science Foundation of China (31801199 to SL and 31801201 to XJ, and 81630059 to HD).

## Author contributions

GJ and FS initiated and supervised the project. GJ and SL designed and constructed the HOPE-SIM system. GJ wrote the software programs. SL performed the cryo-CLEM experiments and cryo-ET work. XJ, XZ and HD performed the cell culturing and vitrification experiments. XJ, GJ and SL performed the cryo-FIB fabrication work. TN, SL, XJ and CQ performed image processing. SL, GJ and FS analyzed the data and wrote the manuscript.

## SUPPLEMENTARY INFORMATION

**Supplementary Note 1. Points to refine the correlation accuracy between the 3D cryo-SIM and 2D cryo-FIB images**.

First, the PT coating protective layer on the surface of the cryo-sample must be thin enough to be able to identify fiducial markers. Second, we recommend performing fine milling with a smaller beam current of 40∼80 pA during the whole cutting process to reduce cutting error caused by the FIB-induced deformation of cryo-lamella. Third, fiducial markers without a complete fluorescence signal or invisible in cryo-FIB images should be preferentially rejected. The standard deviation between cryo-SIM and cryo-FIB images is calculated, which can guide the choice of the number of markers used for correlation. To obtain the best correlation accuracy, there must be sufficient markers around the target. The set of markers with a standard deviation smaller than 200 nm can then be used for further optimization when calculating the correlation.

**Supplementary Figure 1.**
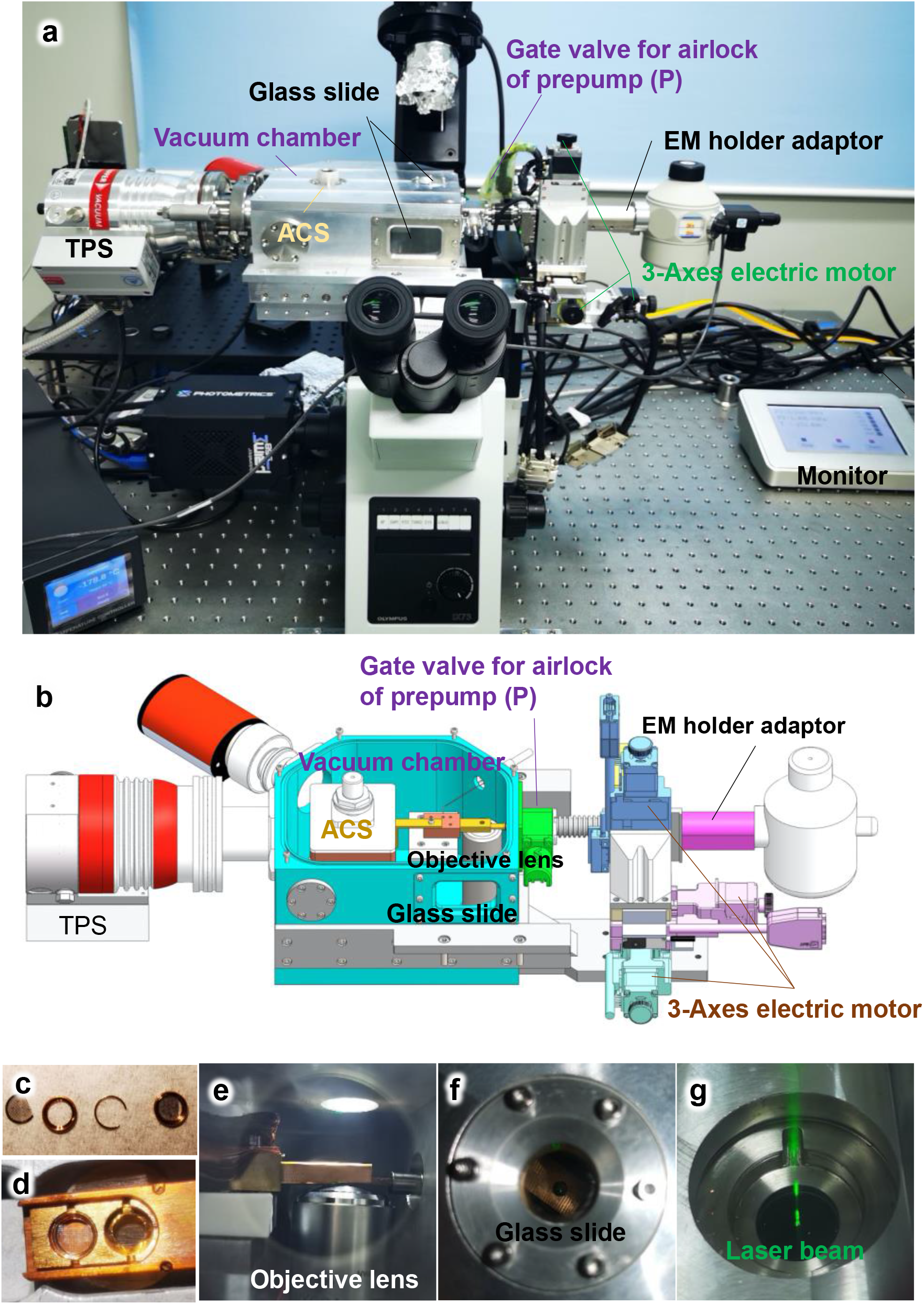
Design and overview of the HOPE-SIM system stage. (a) A real photograph of an Olympus IX73 inverted microscope with the mounted HOPE-SIM stage. TPS, turbo pump system; ACS, anti-contamination system. (b) Section view of the design of the HOPE-SIM stage. Each part of the system is labeled and described. (c) D-shape EM finder grid, AutoGrid and C-clip used in the HOPE-SIM system. (d) Real photograph of a custom multiholder tip mounted with AutoGrid in the transfer workstation. (e) Real-time observation of the objective lens and ACS cryo-box under the working conditions through the observation window. (f) Top glass slide of the HOPE-SIM system. (g) Bottom glass window with the laser beam passing through.

**Supplementary Figure 2.**
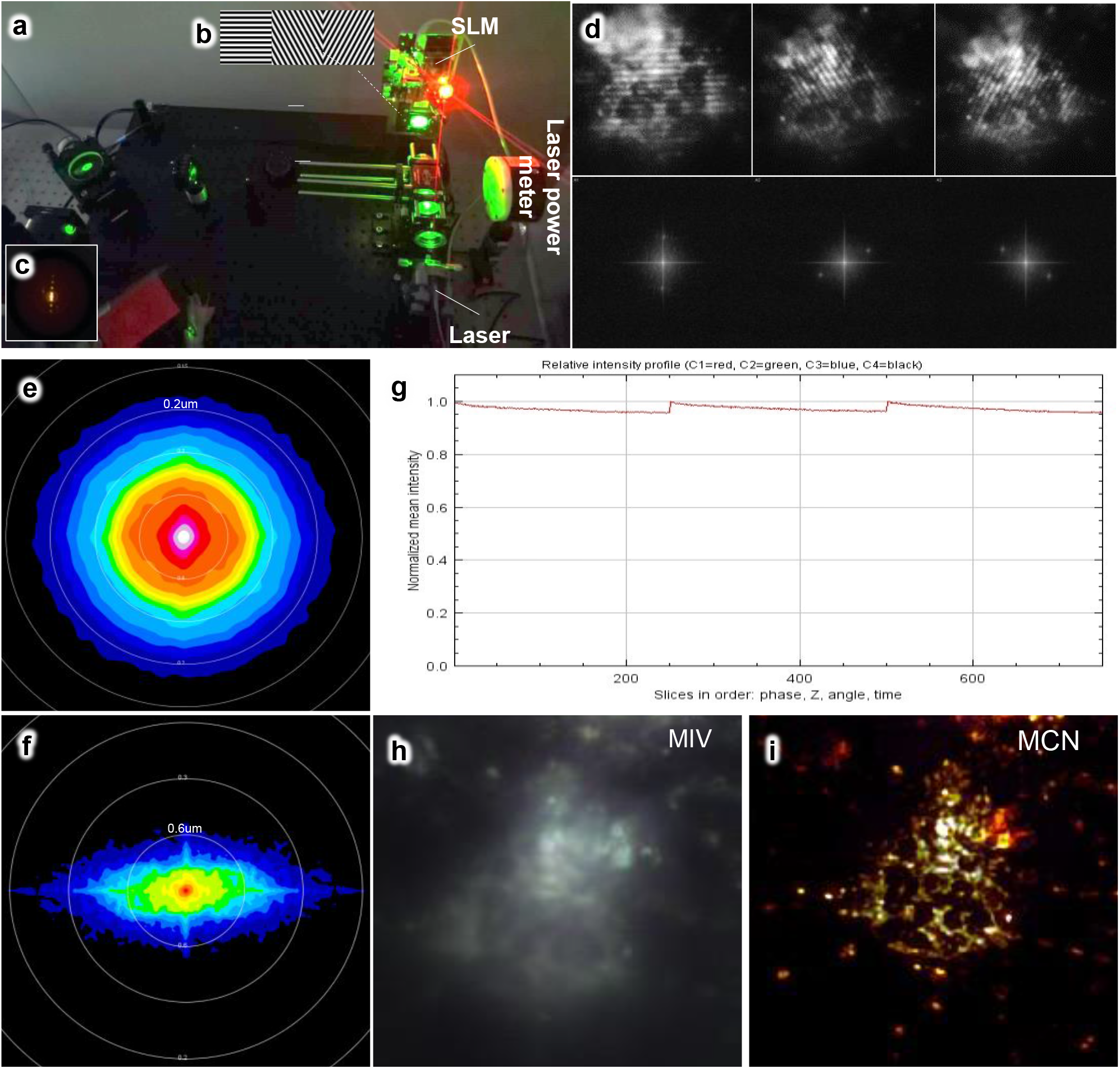
Overview of the HOPE-SIM optical system. (a) A real photograph of the optical system of HOPE-SIM under working conditions. SLM, spatial light modulator. (b) Three patterned illuminations with different rotations generated by SLM. (c) Diffractions of the order 0 and ±1 are focused on the back focal plane of the objective. (d) Representative raw cryo-SIM images (three illumination patterns) of HeLa cells cultured on the grid and stained with MitoTracker Red CMXRos dye. The corresponding Fourier transformation is shown below with the diffraction points on the order of 0 and ±1. (e) Lateral Fourier spectra of the reconstructed image to show the attainable resolution (∼200 nm) in the XY plane. Red, high intensity. Blue, low intensity. (f) Orthogonal Fourier spectra of the reconstructed image to show the attainable resolution (∼500 nm) in the Z direction. (g) Channel intensity profile generated by SIM-Check ^1^ to sequentially show the total fluorescence intensity variation along different phases, z-slices and angles at one time point. (h) Motion and illumination variation (MIV) generated by SIM-Check ^1^ to check the stability of the system during 3D cryo-SIM imaging. The phase-averaged and intensity-normalized images for each angle are merged with different colors. The gray-white appearance of MIV indicates good motion stability during illumination. (i) Modulation contrast-to-noise ratio (MCN) heatmap generated by SIM-Check ^1^ to check the fluorescent signal distribution.

**Supplementary Figure 3.**
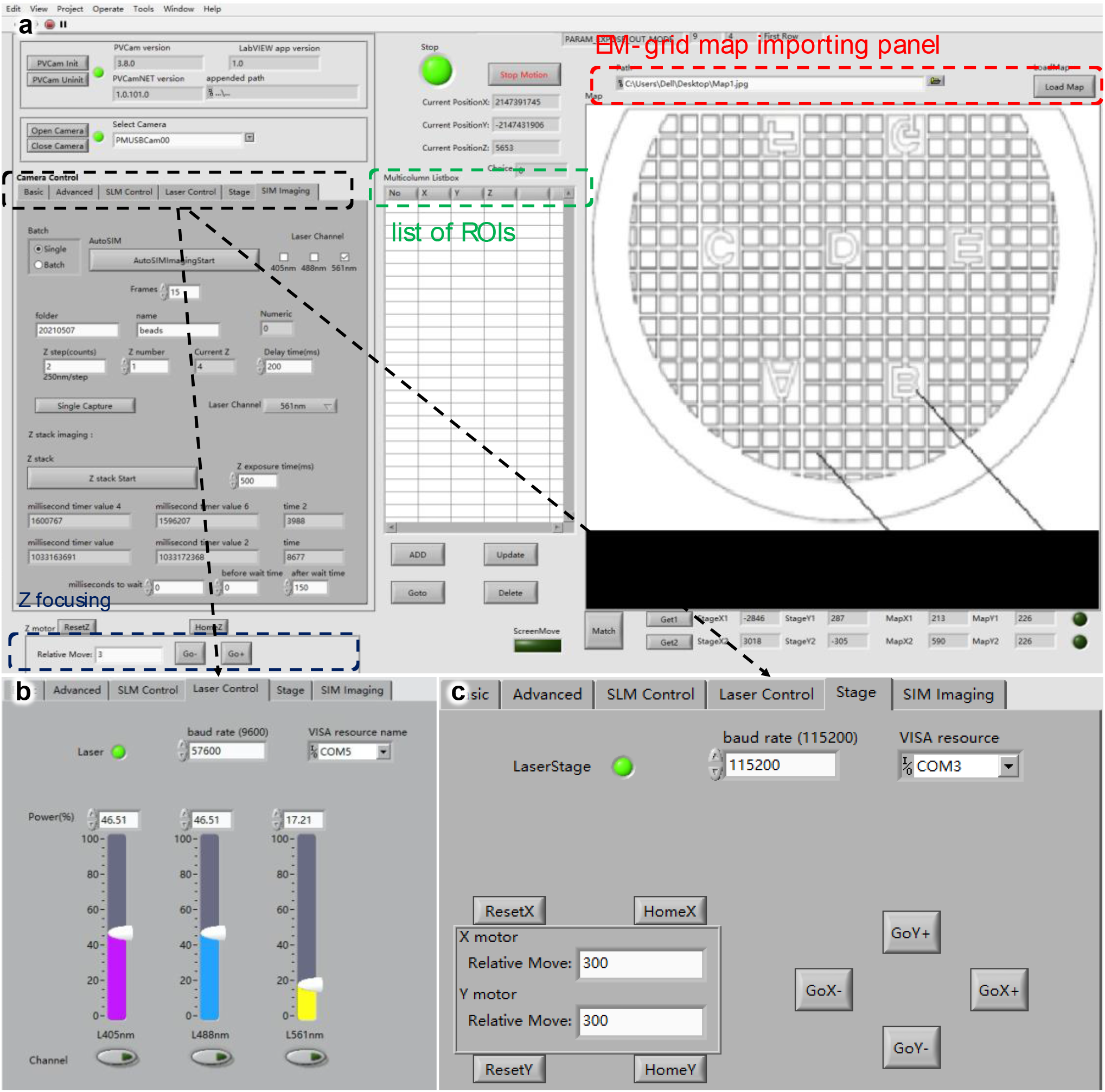
GUI of the HOPE-SIM View software. (a) The main page of HOPE-SIM View that includes the EM-grid map importing panel, list of ROIs panel, z-focus panel and controlling panel. (b) Laser control page. (c) Stage control page.

**Supplementary Figure 4.**
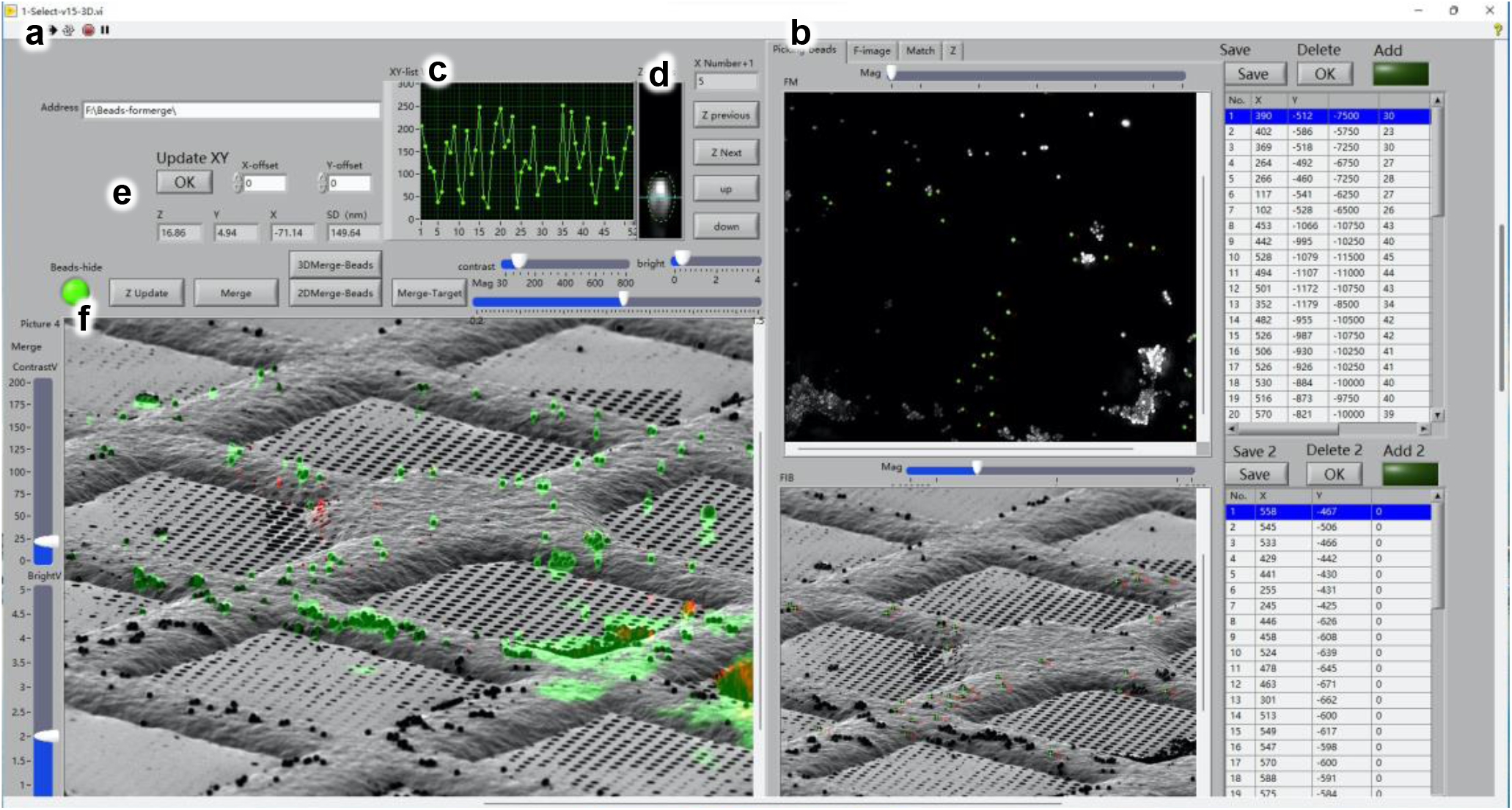
GUI of the 3D View software. The main page of the 3D View software that contains the cryo-FM and cryo-FIB images loader (a), the panels for fiducial marker picking and position listing (b), for plotting the deviations of correlated fiducial markers (c), for fiducial marker Z-height optimization (d), for displaying the correlation parameters (e) and for showing the final merged image after correlation (f).

**Supplementary Figure 5.**
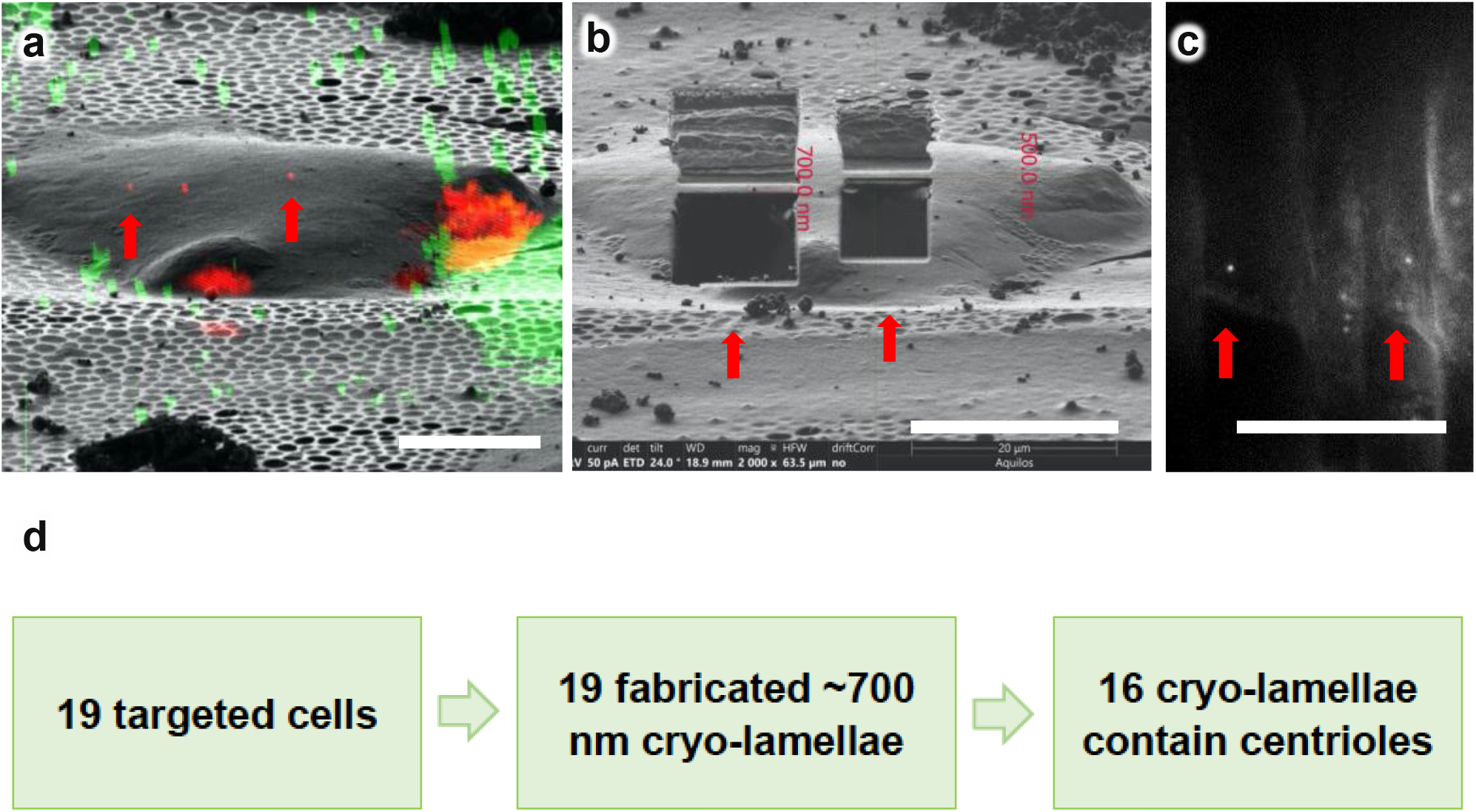
Success rate of targeting centrosomes using the HOPE-SIM cryo-CLEM workflow. (a) 3D correlative image indicates the fluorescence-labeled centrosomes (red) of HeLa cells, which are indicated by red arrows. Green, fluorescent microspheres. (b) Site-specific cryo-FIB fabrication was performed to obtain cryo-lamellae (red arrows) with thicknesses of ∼700 nm. (c) After cryo-FIB milling, the cryo-EM grid containing the cryo-lamellae was transferred back to the HOPE-SIM system and imaged by cryo-SIM. The fluorescence signals of the target centrosomes were captured (red arrows), proving the success of site-specific cryo-FIB using the HOPE-SIM cryo-CLEM workflow (d). See also **Supplementary Data 1**.

**Supplementary Table 1.**
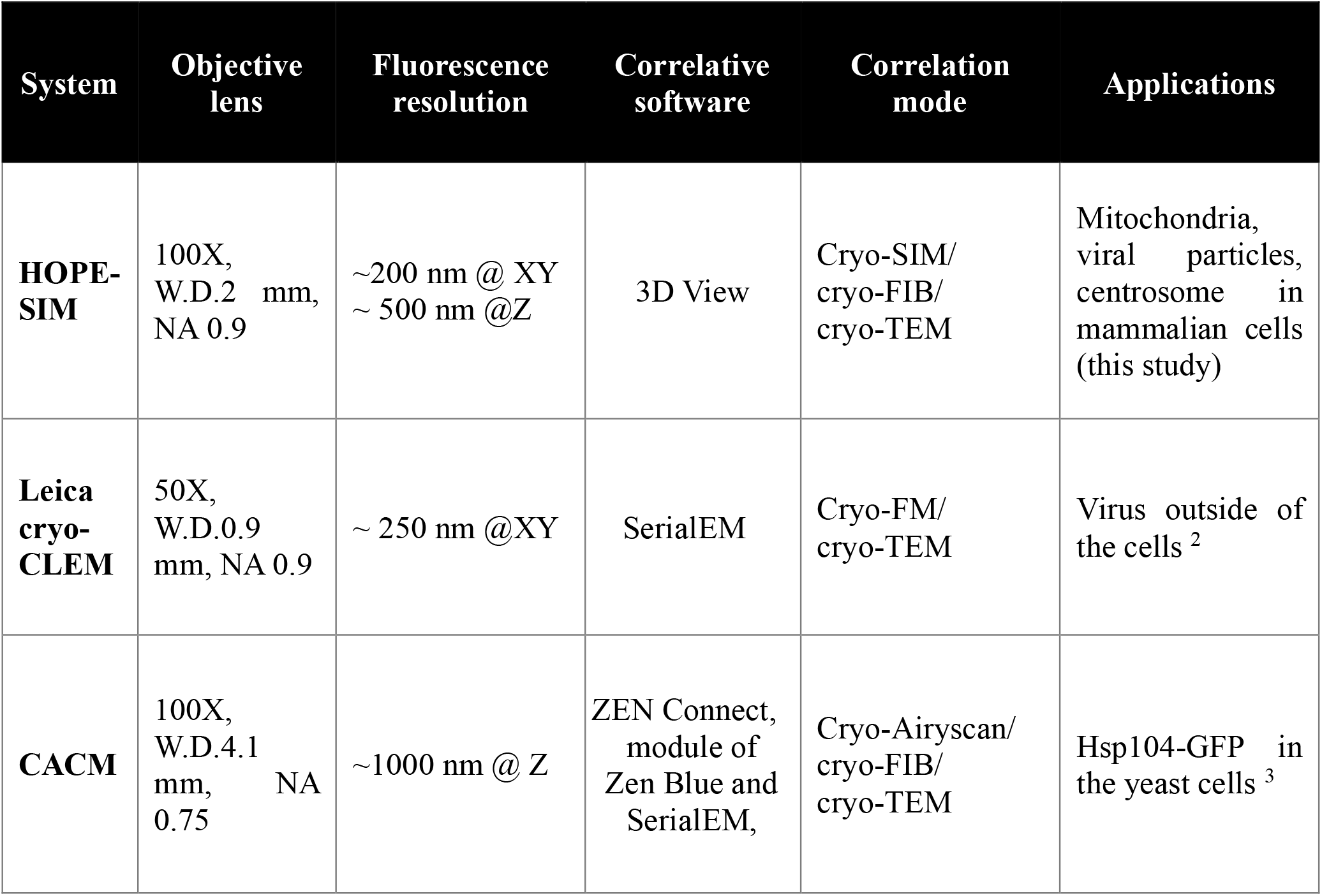
Comparison of characteristics 1 for different cryo-CLEM systems.

**Supplementary Video 1. Design and p 1 rinciple of the HOPE-SIM system**.

**Supplementary Video 2. HOPE-SIM-based cryo-CLEM workflow**.

**Supplementary Video 3. Aligned cryo-ET tilt series of cryo-lamella containing target centrosomes**.

**Supplementary Video 4. Tomogram of cryo-lamella containing target centrosomes**.

**Supplementary Video 5. Another tomogram of thicker cryo-lamella containing target centrosomes**.

**Supplementary Video 6. 3D *in situ* structure of the centriole of HeLa cells rendered on the surface. The triplet tubules are colored yellow, and the internal scaffold is colored pink**.

**Supplementary Data 1. List of merge images of cryo-lamellae with the thickness of ∼700 nm. For each merge image, the fluorescence signal is merged with the wide-field image. The cryo-lamellae are numbered and marked by squared with red for the one containing targeted centrosome and green for the one missing the target**.

